# Intravital imaging of age-related conjunctival lymphatic changes on the ocular surface

**DOI:** 10.64898/2026.06.25.734608

**Authors:** Yujia Yang, Lejun Shen, Erin Luna, Lorissa Zhou, Paula Castillo Hi Espino, Guangyu Li, Lu Chen

## Abstract

**Purpose:** Lymphatic system plays a critical role in fluid regulation and immune response. The conjunctiva of the ocular surface is endowed with a rich lymphatic network, and it provides an ideal site to study lymphatic structure and function. The purpose of this study is to investigate potential morphological and functional changes of conjunctival lymphatics with aging, a time-dependent biological deterioration process.

**Methods:** Young and aged fluorescently labeled transgenic mice of Prox-1, the master control gene for lymphatic determination, were used in the study. For morphological assessment, conjunctival lymphatic vessels were examined in vivo by our advanced live imaging system. For functional analysis, lymphatic drainage efficiencies were measured by fluorescently labeled tracer injection.

**Results:** Compared to young mice, both vascular branching points and intraluminal valves were significantly reduced in conjunctival lymphatic vessels of aged mice. Moreover, lymphatic functional deterioration and drainage deficiencies, such as fluid leakage and reflux, were also detected in the aged condition.

**Conclusions:** Conjunctival lymphatic system undergoes morphological as well as functional changes with aging. Further investigation into this phenomenon may provide novel insights into lymphatic and age-related diseases inside and outside the eye.

## 1. Introduction

The lymphatic network penetrates most tissues in the body and plays critical roles in tissue fluid regulation and immune surveillance. It has been found that lymphatic dysfunction is associated with a wide array of diseases and disorders, which include but are not limited to cancer metastasis, transplant rejection, glaucoma, and inflammatory and immune diseases[1-4]. The current knowledge about aged-related lymphatic changes is rather limited at this stage, which is a topic of our research.

The conjunctiva is a major structure of the ocular surface. It offers an ideal site for lymphatic research due to its anatomical accessibility and transparent nature. This tissue is endowed with a rich network of lymphatic vessels. These vessels express both Prox-1 (prospero homeobox 1) and LYVE-1 (lymphatic vessel endothelial hyaluronan receptor-1) like typical lymphatics in other parts of the body[5-7]. Prox-1 is the master control gene for lymphatic development. Moreover, like typical collecting lymphatic vessels in other organs and tissues, conjunctival lymphatics are equipped with luminal valves for unidirectional fluid flow[8]. The conjunctival lymphatic vessels, therefore, provides a unique opportunity to study the lymphatic system in general.

Unlike blood vessels, lymphatic vessels are not easily visible. In this study, we have employed an advanced live imaging system for in vivo observation of the lymphatic vessels on the ocular surface. The study was performed using a fluorescently labeled transgenic mouse of Prox-1, as we published previously[9, 10]. Taking advantage of these advanced technologies, we were able to visualize and detect age-related morphological as well as functional changes of conjunctival lymphatic vessels in vivo and in real time. This study also establishes a direct and powerful system to explore the general lymphatic system in vivo using the eye as a window.

## 2. Materials and Methods

### 2.1. Animals

As published previously, adult mice of Prox-1-tdTomato (red fluorescent protein) of C57BL/6 background were used in the study[11], in two age groups: 6-month-old and over 24-month-old (n=5/group). All animals were treated according to the ARVO Statement for the Use of Animals in Ophthalmic and Vision Research, and the protocols were approved by the Animal Care and Use Committee, University of California, Berkeley.

### 2.2. Subconjunctival tracer injection

The experiment was performed as we published previously[12]. Mice were anesthetized with isoflurane and topical proparacaine hydrochloride ophthalmic solution (0.5%, Sandoz Inc, Princeton, NJ, USA). Fluorescently labeled tracer (2000kDa, fluorescein-dextran, Invitrogen, USA) was injected into the subconjunctival space with a 33G needle connected to a 10 μL Hamilton syringe (Hamilton, Reno, NV, USA). Tracer uptake was monitored under a fluorescent stereomicroscope connected to a camera and a computer station with imaging software and screen. Antibiotic ointment (Neomycin and polymyxin B sulfates and bacitracin zinc ophthalmic ointment, Bausch & Lomb, Bridgewater, NJ, USA) was applied after the procedure.

### 2.3. Intravital imaging and processing

Intravital imaging was performed using our advanced live imaging system (Zeiss Axio Zoom V.16, Carl Zeiss AG, Gottingen, Germany) as previously reported[6, 9, 12]. Mice were anesthetized with isoflurane and imaged for the superior bulbar conjunctiva. For functional assay after subconjunctival tracer injection, images of three channels (Bright field, Fluorescein, tdTomato) were taken every 30 seconds for 2 minutes. Stack images were processed with Helicon Focus software (Heliconsoft Ltd., http://www.heliconsoft.com) to obtain extended focus.

### 2.4. Statistical Analysis

The results are reported as mean ± SEM. Student t-test was used for morphological assays, two-way ANOVA was used for functional assays. The differences were considered statistically significant when *P* < 0.05 using Prism software (GraphPad, Boston, MA, USA).

## 3. Results

### 3.1. Morphological changes of lymphatic vessels with aging

As shown in Figure 1, a network of lymphatic vessels with luminal valves were detected in the conjunctiva in vivo by the live imaging system. While this network appears denser in the young than the aged mice, further quantitative analysis revealed a significant reduction in both lymphatic branch point and luminal valve number in the aged mice. This finding indicates a structural alteration in the complexity of the lymphatic network with aging.

**Figure 1.**
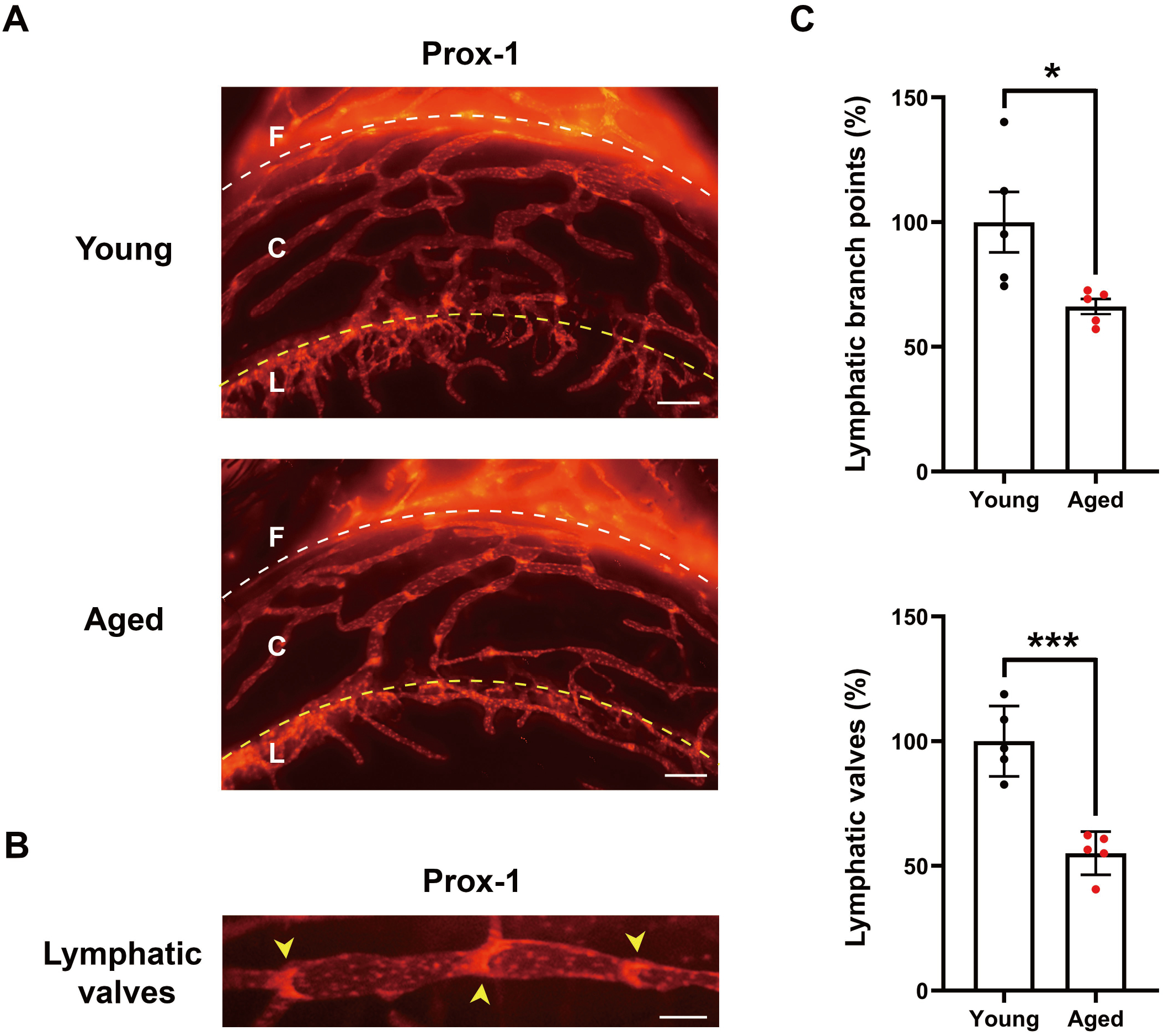
Structural changes of conjunctival lymphatics with aging. A. Representative live images from Prox-1-tdTomato mice showing a denser lymphatic network (red) with more luminal valves in the young condition. F: fornix (white dotted line); C: conjunctiva; L: limbus (yellow dotted line). Scale bars: 100 μm. **B**. Representative live image at higher magnification showing luminal valves (yellow arrowheads) inside the vessels. Scale bars: 50 μm. **C**. Summarized data showing reduced lymphatic vessel branch points and valves in the young than and aged group. * P < 0.05, *** P < 0.005.

### 3.2. Functional deterioration of lymphatic vessels with aging

We next examined the functional changes of conjunctival lymphatics with aging using a lymphatic specific fluorescently labeled tracer, as we reported previously[12]. As demonstrated in Figure 2 in a normal condition, the lymphatic tracer was taken up by the lymphatic vessels after the injection and transported progressively inside the collecting lymphatic channel with time. Compared to the young mice, the aged mice showed a significantly different pattern with unimpeded and faster transportation of the tracer, as presented in Figure 3A. Further quantitative analysis revealed that the tracer traversed a longer distance within the same time post-injection in the aged than the young mice (Figure 3B). This finding indicates a loss of regulation of the lymphatic drainage in the aged condition, corresponding to deteriorated lymphatic and valve function.

**Figure 2.**
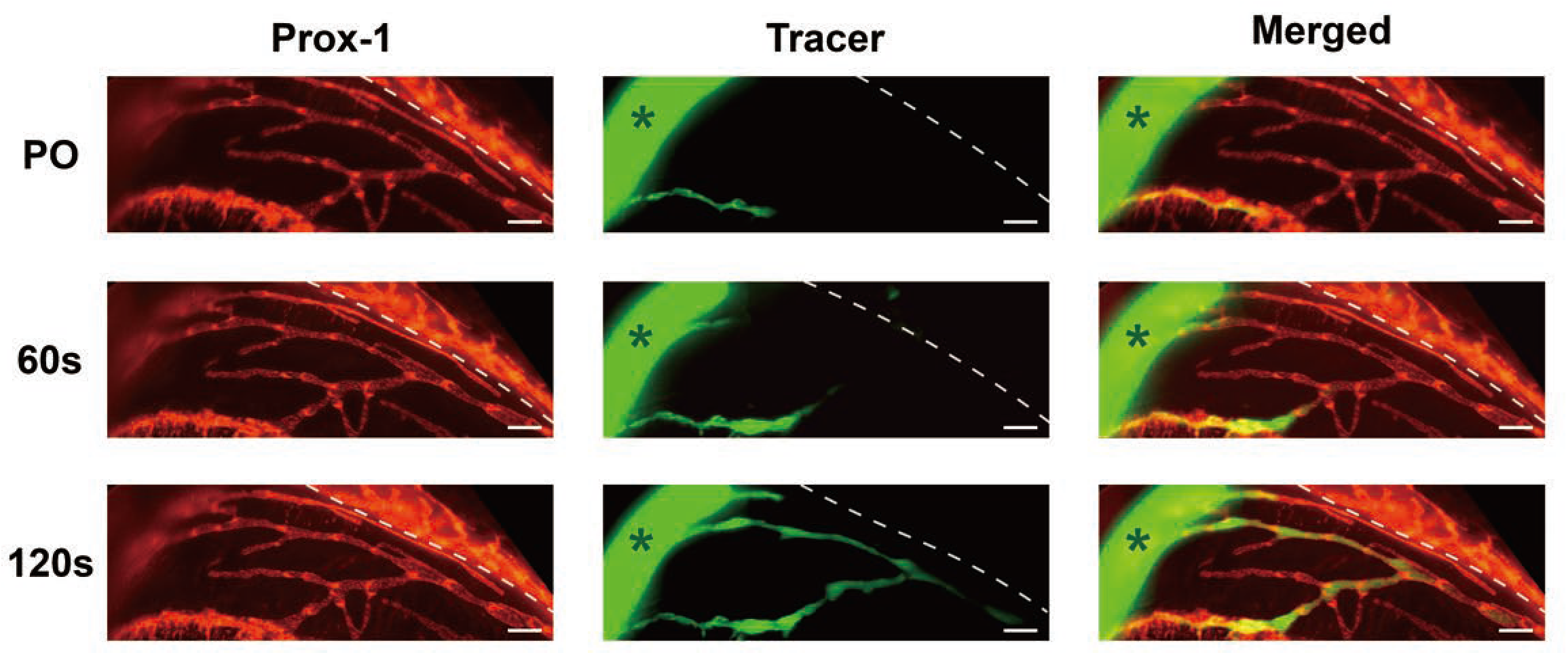
Lymphatic functional analysis by fluorescently labeled tracer injection. Representative live images in a young mouse showing segmental drainage of the lymphatic tracer (green, FITC-dextran) within the lymphatic vessels (red) after subconjunctival injection (asterisk) in a normal condition. PO, post-operation. Scale bar: 100 μm.

**Figure 3.**
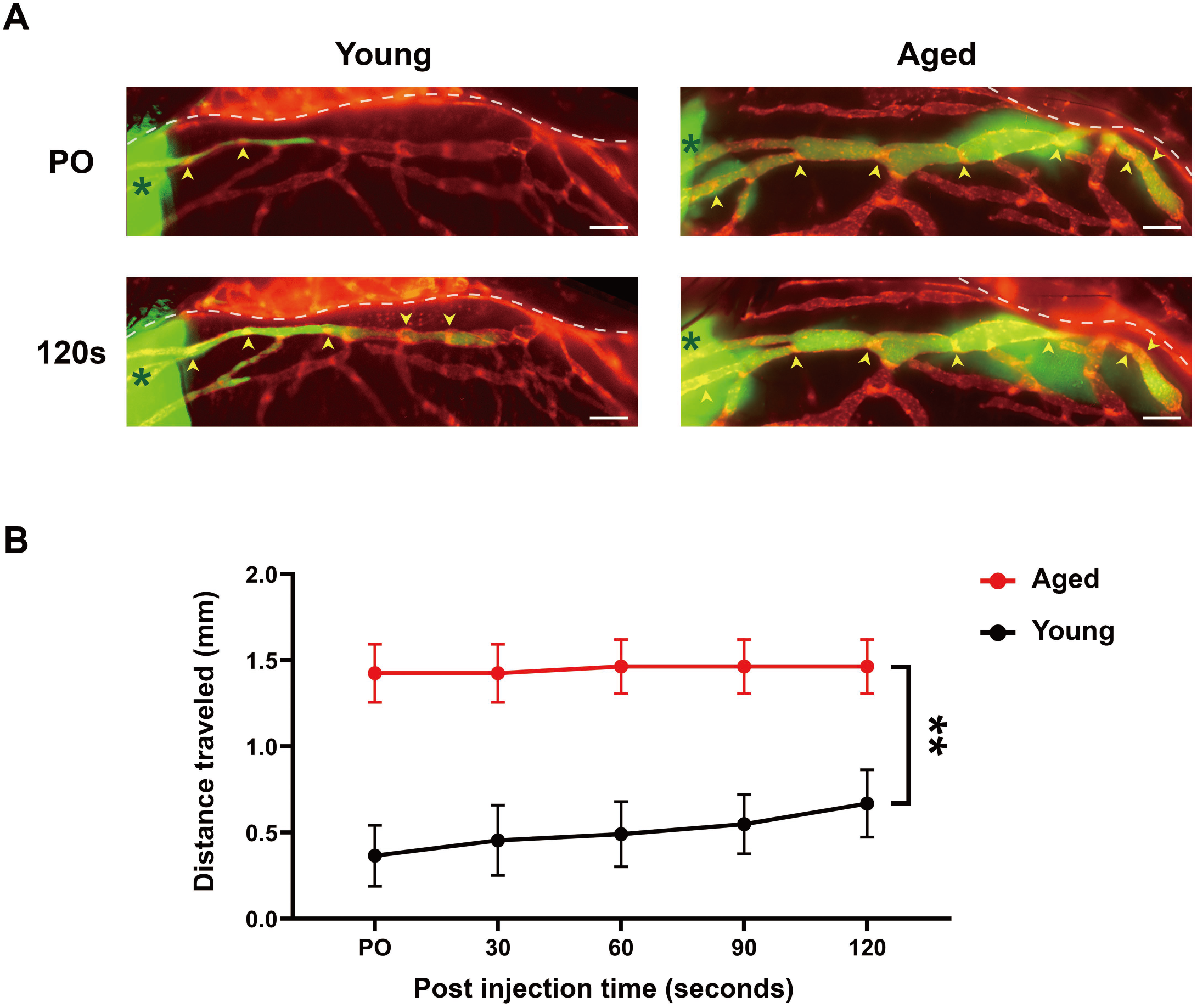
Functional changes of conjunctival lymphatics with aging. A. Representative live images showing defective lymphatic drainage in the aged condition. Arrowheads: valves (red) passed by the lymphatic tracer (green). PO, post-operation. Scale bars: 100 μm. **B**. Summarized data showing faster transportation of the tracer in longer distance in the aged group. ** P < 0.01.

### 3.3. Abnormal lymphatic leakage detected in aged condition

As shown in Figure 4, live imaging of lymphatic tracer tracking also revealed increased lymphatic vessel permeability in the aged group. Compared to the young condition where the tracer was confined within the lymphatic channels, diffused tracer was observed outside the channels in the aged condition, indicating compromised integrity of the conjunctival lymphatics.

**Figure 4.**
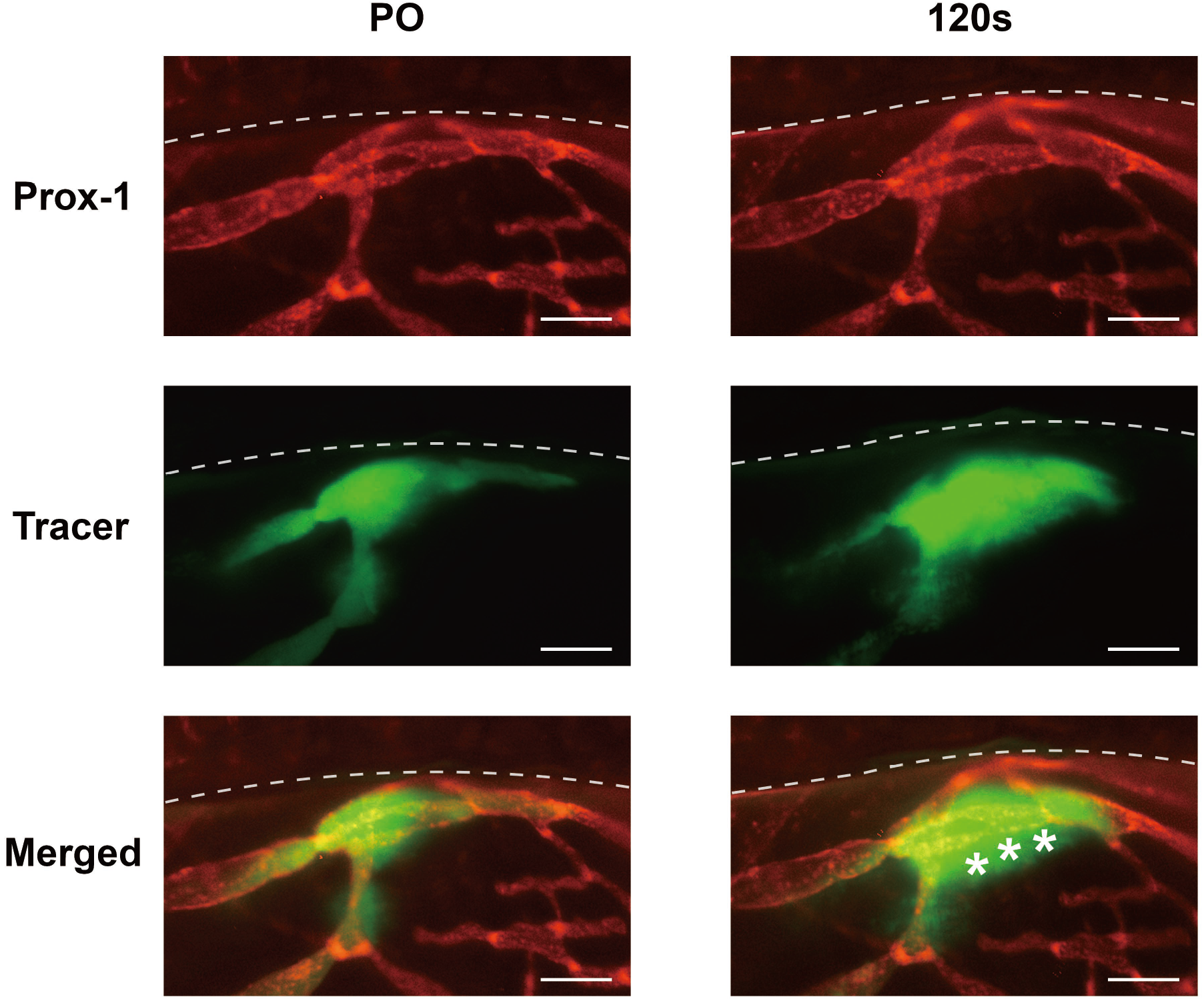
Lymphatic leakage in aged condition. Representative live images showing lymphatic tracer (green) detected outside the lymphatic vessel (red) in an aged mouse, as indicated by the white asterisks. PO, post-operation. Scale bars: 100 μm.

### 3.4. Abnormal lymphatic backflow detected in aged condition

As shown in Figure 5, another functional deficiency, drainage reflux, was also detected in the aged condition. Instead of progressive transportation along the lymphatic channel, the back flow suggests a loss of drainage direction that is normally controlled by unidirectional valves inside the vessels.

**Figure 5.**
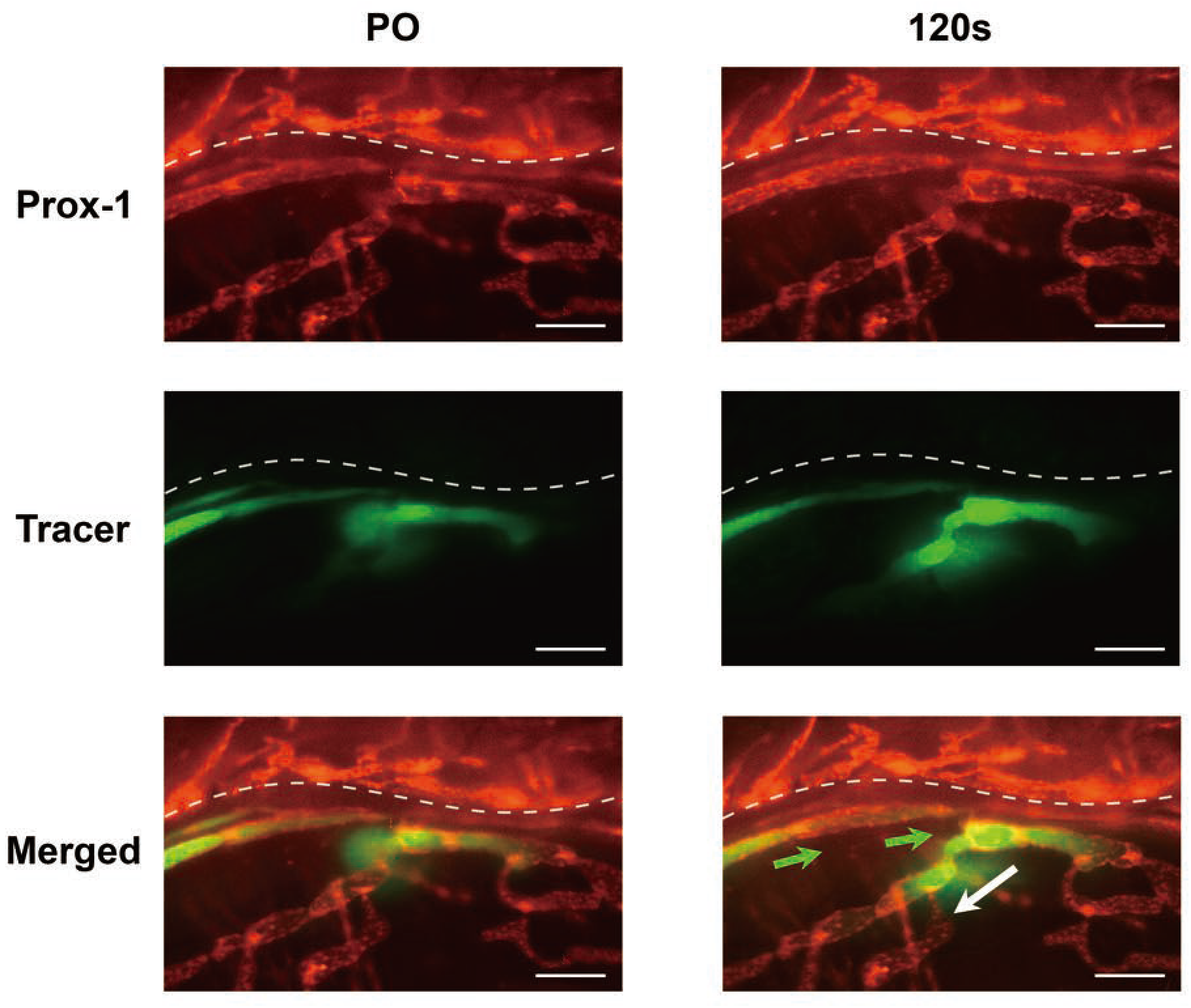
Lymphatic reflux in aged condition. Representative live images showing backflow (indicated by the white arrow) of lymphatic tracer (green) at a branching point. Green arrows: normal drainage direction. PO, post-operation. Scale bars: 100 μm.

## 4. Discussion

In this study, we have shown that aging imposes a serious challenge to the lymphatic system on the ocular surface. Compared to the young condition, both vascular branching points and intraluminal valves were significantly reduced in conjunctival lymphatic vessels of the aged condition. These morphological alterations are coupled with functional deteriorations of the system, as reflected by drainage deficiencies, such as leakage and reflux. Further investigation into these novel phenomena may provide novel insights into age-related diseases in the eye.

Moreover, with this study, we have demonstrated that the conjunctiva can serve as an ideal site to study lymphatic biology and function. Sitting on the ocular surface as a transparent tissue, it allows direct and noninvasive observation of the lymphatic network in vivo and in real time.

Yet to be explored, it is possible to employ this platform to investigate the interplay between the lymphatic network with other cellular and structural components in the environment, such as immune cells and blood vessels, or to assess other biological or pathological processes and the therapeutic effects of an intervention. Knowledge obtained from conjunctival lymphatic research can be readily translated to other tissues and organs harboring the lymphatic network but not easily accessible.

This study provides new evidence supporting lymphatic dysfunction as a feature of aging. Our results are consistent with other reports outside the eye, such as the skin[13], mesentery[14], and meninges[15, 16]. Taken together, these data have identified lymphatic dysfunction as a general parameter and biological marker for aging across different tissues and organs. This functional change may lead to further pathological responses and tissue deteriorations, which warrants future investigation. Besides the lymphatic system alone, age-related impairment of the glymphatic/lymphatic system is also identified in the meninge, and reduced clearance of metabolic waste has been considered as a contributing factor to the pathogenesis of neurodegenerative diseases, such as Alzheimer’s disease[16, 17]. It is anticipated that future studies on the fluid transportation systems, lymphatics and/or glymphatics, may provide novel mechanisms and therapeutic approaches for a broad spectrum of diseases occurring inside and outside the eye.

## Acknowledgements

This work is supported in part by research grants from National Institutes of Health and University of California at Berkeley. The authors thank Young K. Hong from University of Southern California and the Mutant Mouse Regional Resource Centers (MMRRC) for providing the founder Prox-1 transgenic mice.

